# Hydroxyapatite pellets as versatile model surfaces for systematic studies on enamel

**DOI:** 10.1101/2021.01.11.426207

**Authors:** Johannes Mischo, Thomas Faidt, Ryan B. McMillan, Johanna Dudek, Gubesh Gunaratnam, Pardis Bayenat, Anne Holtsch, Christian Spengler, Frank Müller, Hendrik Hähl, Markus Bischoff, Matthias Hannig, Karin Jacobs

## Abstract

Research into materials for medical application draws inspiration from naturally occurring or synthesized surfaces, just like many other research directions. For medical application of materials, particular attention has to be paid to biocompatibility, osseointegration and bacterial adhesion behavior. To understand their properties and behavior, experimental studies with natural materials such as teeth are strongly required. The results, however, may be highly case-dependent because natural surfaces have the disadvantage of being subject to wide variations, for instance in their chemical composition, structure, morphology, roughness, and porosity. A synthetic surface which mimics enamel in its performance with respect to bacterial adhesion and biocompatibility would, therefore, facilitate systematic studies much better. In this study, we discuss the possibility of using hydroxyapatite (HAp) pellets to simulate the surfaces of teeth and show the possibility and limitations of using a model surface. We performed single-cell force spectroscopy with single *Staphylococcus aureus* cells to measure adhesion-related parameters such as adhesion force and rupture length of adhesins binding to HAp and enamel. We also examine the influence of blood plasma and saliva on the adhesion properties of *S. aureus*. The results of these measurements are matched to water wettability, elemental composition of the samples and the change in the macromolecules adsorbed over time. We found that the adhesion properties of *S. aureus* were similar on both samples under all conditions: Significant decreases in adhesion strength were found equally in the presence of saliva or blood plasma on both surfaces. We therefore conclude that HAp pellets are a good alternative for natural dental material. This is especially true when slight variations in the physicochemical properties of the natural materials may affect the experimental series.

## Introduction

Hydroxyapatite (HAp, Ca_10_(PO_4_)_6_(OH)_2_) is the main mineral component of human enamel as well as of bones [1]. As an integral and structural part of the body, research into HAp as a biomaterial, its synthesis, application development and improvements has progressed over the last decades [1]: for instance, HAp-based cements are readily available for use, robust HAp compounds with high fracture toughness and wear resistance have been developed and porous HAp scaffolds for bone regeneration have been proposed [1–3]. In most modern medical application, natural HAp-based components like bone or tooth are still most often mended with implant materials, such as titanium in artificial hip joints or screws [4, 5]. Despite advancements in mechanical aspects, osseointegration, corrosion resistance and metal ion release of medical implants over the last years, biomaterial-centered infections are still quite common and often lead to severe medical complications such as prosthetic implant failures or aseptic loosening [6–9]. These are commonly associated with bacterial biofilm formation on the implant surface and often require removal of the implant material for curing [10]. For dental implants for instance, a 2015 study of over one thousand dental implants shows that up to 10 % of patients receiving dental implants suffer from postoperative infections, two-thirds of which must have their implant removed [11]. To improve the success of implantation, the prevention of biofilm formation on the implant surface is integral [12]. Research into bacterial biofilm formation is, however, in most cases either carried out on highly artificial laboratory surfaces, such as silicon wafers or glass slides, or on natural samples, that are subject to severe sample-to-sample changes due to external factors like age, material composition and morphology [13–16]. We propose systematic studies of factors influencing biofilm formation, using surfaces that offer both the verisimilitude of a natural material and the advantages of reproducibility and well-defined material properties, such as surface topography.

A material that meets the requirements of closeness to natural materials such as enamel has to mimic both the biocompatibility and characteristics concerning bacterial biofilms, such a natural material has. Bacterial infections start with the adhesion of single, planktonic bacteria to a surface. The bacteria then start to grow into microorganism consortia, embedded in an extracellular matrix in which the bacteria are protected from host defense mechanisms and antibacterial therapy [17, 18]. A common approach is to prevent or hinder the bacterial adhesion process as the first step of bacterial biofilm formation [19]. However, many of the coatings or substances found to be antibacterial in the laboratory, such as silver, lack osseointegration or mammalian cell growth and attachment is simultaneously suppressed when applied [12]. In this paper, we evaluate HAp as a basis for further systematic research, as HAp is a mineral synthesized by mammals and is known for its biocompatibility and osseointegration [20–23]. We compare surface properties of artificially synthesized HAp pellets to natural bovine enamel, investigate single-bacterium adhesion and determine the underlying adhesion forces and ruptures lengths.

Single-cell force spectroscopy (SCFS) with an atomic force microscope has proven to be an ideal method to determine the adhesion parameters of bacterial cell wall macromolecules forming interactions with the surface [24–26]. For the purpose of this study, we have chosen the opportunistic pathogen *Staphylococcus aureus*, as it forms clinically relevant biofilms [27], and is a common cause of implant failures and inflammation in the oral cavity [28–30] and beyond [10].

It is known that the surface chemistry, hydrophobicity and surface charge [31, 32] or functional groups deposited on top of a surface influence the bacterial adhesion process. Such functional groups could be for instance proteins, such as silk proteins [33, 34], or bodily fluids such as blood plasma [35, 36] or salivary macromolecules [37, 38, 14]. For the purpose of this paper, we therefore use well-characterized samples and also examine the influence of conditioning films consisting exclusively of the macromolecules present in either saliva or blood plasma because any material in the body is inevitably in contact with host fluids. For example, it has been shown that a salivary macromolecule film, termed pellicle, forms within seconds after a surface is brought into contact with saliva [39]. The pellicle reaches a thickness of around 7 nm within three minutes and is free from bacteria at this early stage [40, 41]. We also incubate bacteria in saliva to mimic the natural case, in which a bacterium comes into contact with the surfaces we chose from the oral cavity setting. Regardless of whether HAp is suitable as a model surface of dental enamel and thus also for further systematic studies, we expect a reduction of bacterial adhesion strength in the presence of biomolecules of body fluids based on existing literature [36]. A well-characterized, artificially synthesized surface that mimics enamel in its performance with respect to bacterial adhesion and biocompatibility would be an excellent basis for further, systematic studies on parameters influencing these properties.

## Materials and Methods

### Bacteria and bacterial probes

For each experiment, *Staphylococcus aureus* strain SA113 was freshly cultured as follows: The bacteria were inoculated and grown from a deep-frozen glycerol stock on Tryptic Soy Agar Plates with 5 % sheep blood (Becton Dickinson, Heidelberg, Germany) at 37°C for 24 h. A discrete colony was resuspended in Tryptic Soy Broth (TSB, Becton Dickinson) at a culture to flask volume of 1:10 and cultivated at 37°C and 150 rpm for 16 h. To obtain cells from the exponential growth phase, the bacterial solution was diluted by 1:100 in fresh TSB and incubated for another 2.5 h at the same settings, resulting in a bacterial solution with an Optical Density (OD 600) of 0.5. We removed the debris and extracellular material by washing 1 ml of bacterial solution three times using 1 ml of phosphate buffered saline (PBS, pH 7.4, Carl Roth GmbH, Karlsruhe, Germany) as replacement supernatant after centrifuging at 20 000 rcf. The bacterial solution was set to an OD 600 of 0.1. Before each AFM tip functionalization, the bacteria were vortexed to disrupt bacterial aggregates and subsequently diluted 1:100 in PBS. A drop of the diluted bacterial solution was spotted in a petri dish, and a single bacterium was then attached to a calibrated tipless AFM cantilever (MLCT-O10 E, Bruker-Nano, Santa Barbara, US-CA) via dopamine using the technique described by Thewes et al. [42]. This functionalization was controlled optically with an inverted microscope before and after the measurement.

### Human samples

The saliva was donated by five volunteers, both male and female, over 18 years of age, with good oral health. Human blood plasma (BP) was obtained from male healthy volunteers older than 18 years. All subjects gave their informed written consent to participate in this study. Pellicle collection and blood plasma protocols were approved by the medical ethic committee of the Medical Association of Saarland, Germany (code numbers 39/20 and 238/03 2016).

Saliva samples were obtained 1.5 h after tooth brushing with toothpaste (dentalux COMPLEX3 Mint Fresh, DENTAL-Kosmetik GmbH, Dresden, Germany). In between the donors brushed their teeth once without tooth paste one hour after first brushing, and refrained from eating and drinking (except for still water) for the whole time. The saliva obtained was centrifuged at room temperature and 25 000 rcf for 10 min in Falcon tubes (Corning Inc., Corning, US-NY). The supernatant was then transferred to fresh tubes and the process was repeated once. The remaining saliva from all five participants was mixed, aliquoted and stored at –20 °C.

The Human BP was derived from freshly drawn blood and centrifuged at 6 000 rcf in S-Monovette lithium-heparin blood collection tubes (Sarstedt, Nümbrecht, Germany) for 2 min. The plasma was transferred to a fresh reaction tube and centrifuged one more time under the same conditions to remove any remaining cell material. The blood plasma was stored in a fresh reaction tube at –80 °C until usage.

### Sample preparation

The hydroxyapatite (HAp) samples were made from compressed and sintered HAp powder (Sigma Aldrich, Steinheim, Germany) according to the protocol described by Zeitz and Faidt et al. [43]. Before usage, the HAp samples were polished, using abrasive paper (SiC, Struers, Willich, Germany) with decreasing coarseness and polishing solution (MSY 0-0.03, Microdiamant, Lengwil, Switzerland: 30 nm diamond particle solution). The debris from polishing was removed by etching in a sodium acetate buffer (pH 4.5) for 7 s and subsequent sonication in ultrapure water (TKA-GenPure, Thermo Fischer Scientific, Waltham, US-MA). The HAp samples used have the same crystal structure, chemical composition and surface roughness as specified by Zeitz and Faidt et al. [43].

Throughout this study, a single piece of enamel cut from the vestibular surface of a bovine incisor tooth was used. Similar to the HAp sample, the enamel was polished in several steps before usage except for the polishing solution, where a suspension of 40 nm sized colloidal silica particles (OP-S, Struers, Ballerup, Denmark, rebranded: now OP-U) was used. Residues were removed in an ethanol ultrasonic bath.

The generation of conditioning films on both samples was carried out following these procedures: For a BP coating, the samples were incubated for 30 min at 37 °C under humid conditions to prevent the biological coating from drying. For salivary pellicle formation, saliva was applied onto the surfaces and incubated for 3 min at room temperature [40]. For both coatings, the surfaces were washed with PBS and thereafter kept in fresh PBS. Bacteria, which were exposed to saliva before measurement, were incubated in their immobilized state on the tip of a cantilever for 3 min in 25 μl of saliva at room temperature and the whole cantilever was washed afterwards in PBS [38].

### Sample characterization

Surface topography of all samples used was acquired by atomic force microscopy (FastScan Bio, Bruker-Nano, Santa Barbara, US-CA). The instrument was operated with Olympus OMCL-AC160TS probes (Tokyo, Japan) in tapping mode. Roughness values were calculated from 3D scans of 1 μm x 1 μm (512 x 512 pixels) regions captured at a scan rate of 0.1 Hz. Tilt and scan line height jumps were removed (routines “PlaneFit 1^st^ order and “Flatten 0^th^ order”) using the Nanoscope Analysis 1.9 (Bruker Nano, Santa Barbara, US-CA) software.

The wettability of all surfaces was evaluated by water contact angles in fresh ultrapure water using the sessile drop setting on an OCA 25 instrument (Dataphysics, Filderstadt, Germany). To maintain a wet environment for the coated surfaces, all samples were measured in a water bath with air bubbles pressed onto them to determine advancing and receding contact angles.

X-ray photoelectron spectroscopy (XPS) was performed with non-monochromatized Al-K α excitation (ħω = 1486.6 eV) using an ESCALAB MKII spectrometer (Vacuum Generators, Hastings, UK, base pressure approx. 10 ^−10^ mbar). The spectra were normalized by the photoemission cross-sections, as proposed by Yeh and Lindau and adjusted with a Shirley background [44, 45].

To determine the molecular weight of the macromolecules of the conditioning films we performed sodium dodecyl sulfate polyacrylamide gel electrophoresis (SDS-PAGE) and Coomassie-staining as previously described by Trautmann et al. at two different ages of the films [41]. The specimens consisting exclusively of enamel with a total surface area of 8 cm^2^ were purified with 3 % NaOCl, washed with water, ultrasonicated in 70 % isopropanol, and air-dried before exposure to either saliva or PBS for 3 min at room temperature. Non-adsorbed material was removed with 20 ml ultrapure water from a pressure cylinder (Buerkle GmbH, Bad Bellingen, Germany). Then the samples were either directly treated with elution buffer to elute the adsorbed macromolecules or were incubated in PBS at room temperature and the elution was performed 3.5 h later. The elution, subsequent precipitation and final preparation for SDS-PAGE and Coomassie-staining were conducted [41].

### Measurement of adhesion forces

With the bacterial probes described above, force-distance measurements on all surfaces were performed with a Bioscope Catalyst AFM (Bruker-Nano, Santa Barbara, US-CA). The force trigger was set to 300 pN and the lateral distance between two force-distance measurement points was set to 1 μm. The contact time of the bacterium and the surfaces was tested at both the minimal possible time of a few milliseconds (called 0 s surface delay) [46] and 5 s surface delay, during which the force trigger was kept constant.

For the purpose of this paper, we focus on the data with 5 s surface delay and attach measurements with 0 s delay in the supporting information (SI fig. S1). From a force-distance curve it is possible to obtain values for the adhesion force (see fig. 1a) and rupture length and further detailed information on adhesin-specific unbinding events [46, 32], of which the latter ones are not subject to this paper. For every bacterial cell on each surface, a set of 50 force-distance curves in different spots on the surface for each contact time was recorded and evaluated (see fig. 1b). Throughout the measurement of a single bacterium, no significant changes in the force curves such as decreasing adhesion force were observed. Our measurement procedure was repeated with several cells per surface and the data of all cells were combined (see fig. 1c). The order in which the surfaces were probed after each other had been alternated and had no significant effect.

**Figure 1.**
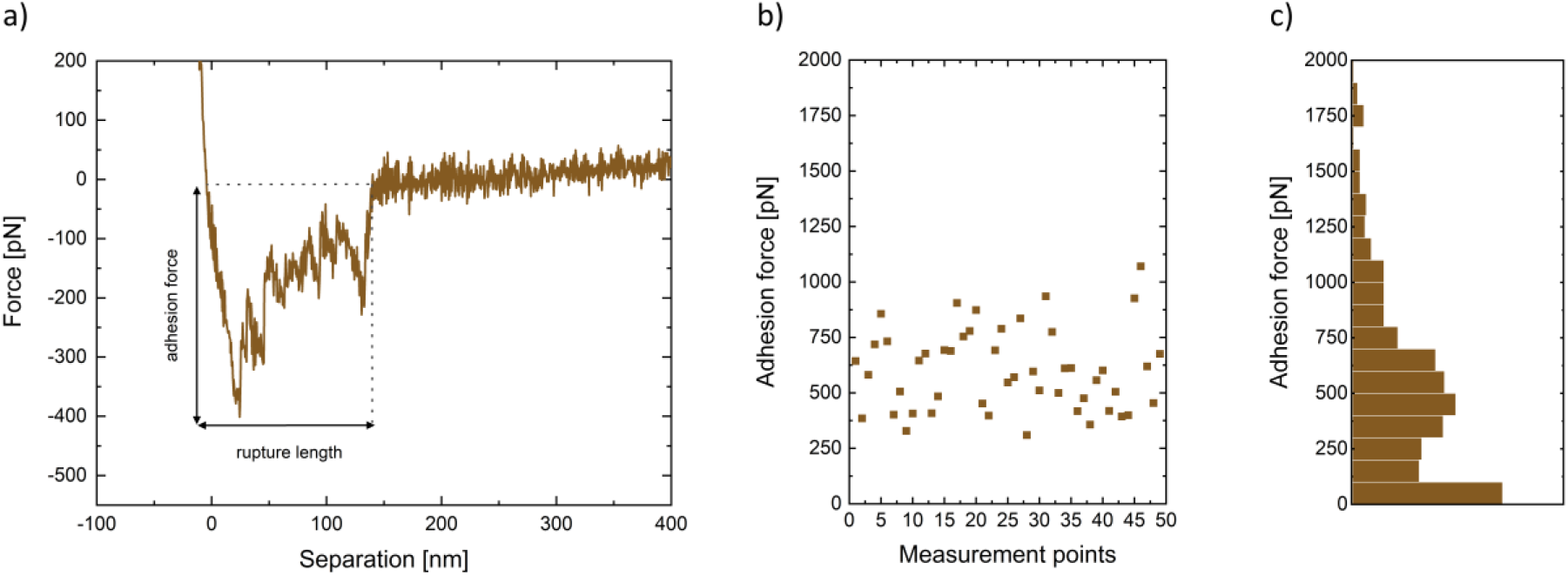
Systematic data acquisition process from a) exemplary force-distance curve of a single bacterium on saliva-coated HAp, to b) the same bacterium was pressed on different places on the surface and the adhesion force was recorded, and c) distribution of several bacterial cell’s adhesion forces represented in a histogram.

### Statistical analysis

Statistical analysis of all data distributions of all coating conditions was conducted, using the Mann-Whitney-U test implemented in the OriginPro2019b software (OriginLab, Northampton, US-MA). The significance levels obtained are presented above the measurement data in figure 3 by asterisks in increasing levels of significance: “n.s.”: not significant; “*”: p < .05; “**”: p < .01; “***”: p < .001 (highest significance).

## Results and Discussions

### Changes in the conditioning film

Single-cell force spectroscopy measurements are performed over hours, during which the adsorbed conditioning films are kept in buffer solution. To monitor the changes such biological surface coatings undergo during our measurement time, we looked at changes in the composition of macromolecules on the surfaces.

Our SDS-PAGE eluates of saliva and BP conditioning films show that the molecular weight of macromolecules adsorbed on the surface does not change markedly between start of the experiments and after 3.5 h in PBS (see SI fig. S2). No distinct statement of the amount of macromolecules adsorbed can be reached as small differences in color depth in the lanes could also be attributed to slight variations of the staining agents. Even if we consider the small decrease in color depth as a reduction of adsorbed macromolecules, no correlations between adhesion force and adsorbed amount or rupture length and adsorbed amount were found. This is probably due to the fact that variations in adhesion forces and rupture lengths are be quite large between individual bacteria [19, 48, 49, 38] and thus much greater than the effect expected from the conditioning film.

### Sample characterization

XPS reveals that HAp and enamel are very similar in chemical composition, except that HAp does not contain any magnesium, sodium or fluorine (see SI fig. S3). The root mean square roughness (RMS) of the hydroxyapatite is 0.46 (0.44) nm and of enamel is 1.27 (0.29) nm. In the literature, increased surface roughness has been found both to increase and decrease adhesion [31, 50–53]. To exclude the influence of roughness, all surfaces are polished to be only rough in a comparable nanometer range.

Imperfections such as deeper grooves from coarser polishing steps or natural crystal boundaries or cracks in the material have no influence on SCFS, as they are negligible due to statistical variation of the cantilever position on the surface (50 positions were probed with each bacterium). Furthermore, the small contact area of *S. aureus* (150 nm to 350 nm radius) [54] makes incidental measurements in such spots even less probable.

The imperfections of the surfaces, however, have an influence on the contact angle measurement in form of pinning and a resulting contact angle hysteresis [55]. For the purpose of this paper, regions with extreme pinning have therefore been omitted during measurement. For the data capture, we used air bubbles to determine the optical contact angle as our samples, both with and without conditioning films (see table 1), have proven to be so hydrophilic that the contact angles of water droplets could not be determined optically. The contact angle is a typical surface property measured in combination with adhesion. It has been shown that the hydrophobicity greatly influences the adhesion of bacteria to a surface in both ways, but in general bacteria adhere better to hydrophobic surfaces [29, 32, 56, 57]. The contact angle is amongst other properties influenced by surface roughness which often varies between surfaces. The influence of surface roughness on the contact angle could therefore be a reason for different findings and should therefore not be underestimated [31, 53]. Lorenzetti et al., for example, showed that the influence of roughness on bacterial adhesion on hydrophilic titanium is much bigger than the influence of contact angle changes on the same surface [31]. On the comparably smooth surfaces used in this study, we expect no greater influence of roughness. On both HAp and enamel, a conditioning protein film leads to a decrease in the mean adhesion force by 10 to 18 % and in the contact angle decreases by up to 70 % compared to the uncoated samples (see fig. 2). Whether the conditioning film on enamel was formed from BP or saliva has, however, only a small influence on the adhesion force. On HAp, we even observe no change in contact angle while the adhesion force decreases for saliva. We therefore conclude that on these surfaces the presence of a conditioning film influences the adhesion force and not the contact angle, which is in accordance with other studies [31, 58].

**Table 1.** Advancing water contact angle (Adv. CA) and the contact angle hysteresis on all surfaces with and without different conditioning films used, averaged over three independent measurements.

|  | No conditioning film |  | Salivary conditioning film |  | BP conditioning film |  |
| --- | --- | --- | --- | --- | --- | --- |
|  | HAp | Enamel | HAp | Enamel | HAp | Enamel |
| Adv. CA [°] | 42.4 | 22.3 | 18.7 | 22.3 | 12.2 | 16.0 |
| Std. Dev. | (4.1) | (2.3) | (3.5) | (2.0) | (3.5) | (4.2) |
| Hysteresis [°] | 14.9 | 0.3 | 0.8 | 0.9 | 0.3 | 1.4 |

**Figure 2.**
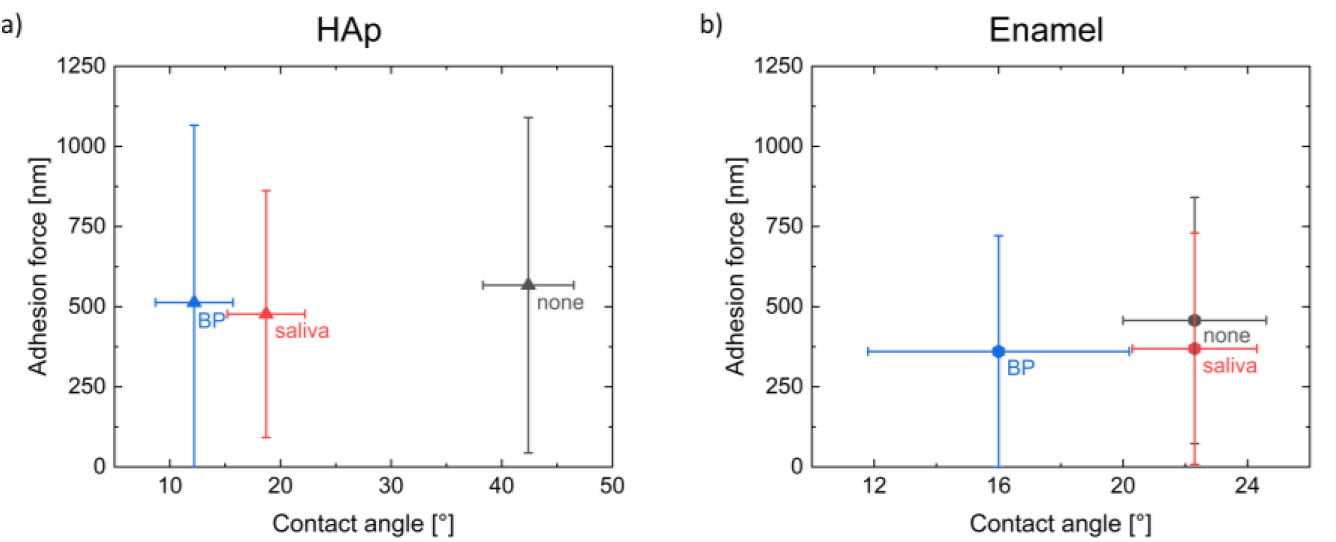
Mean adhesion force including standard deviation versus contact angle including hysteresis on a) HAp and b) enamel for different conditioning films.

### Adhesion force and rupture length

Overall, 146 individual bacteria have been tested in five different combinations of bodily fluid incubations on both the surface and the bacterium (see fig. 3a). The number of measurements per surface and condition is almost evenly distributed, providing good statistics for each surface/condition combination, as individual bacterial cells under the same conditions show larger variations in their adhesion behavior on hydrophilic surfaces [32, 48, 49]. In figure 3, each combination is represented by a separate column containing the data in form of a boxplot and a histogram next to it. For the adhesion measurements, we have chosen data binning in 100 pN steps for the adhesion force measurements and 20 nm steps for the rupture length measurements. In the lowest bin, the majority of data captured is close to or not distinguishable from the instrument’s noise and, therefore, the lowest bin of the adhesion force is considered as no adhesion for the purpose of this paper. To quantify this, the percentage of measurements below 100 pN is indicated next to the lowest bin (see fig. 3a). As the number of measurements observed below 100 pN is quite high for some combinations and thus would dwarf all other bins if they were plotted to scale, all distributions of adhesion forces > 100 pN were scaled to their respective maximum per column for better visibility (excluding the 0 – 100 pN bin). The correlations between the protein combinations from the Whitney-Mann-U test are given above the data in the format “*/*”. The correlations given before the slashes are calculated excluding the lowest bin, while the values after the slashes including the 0 – 100 pN values.

**Figure 3.**
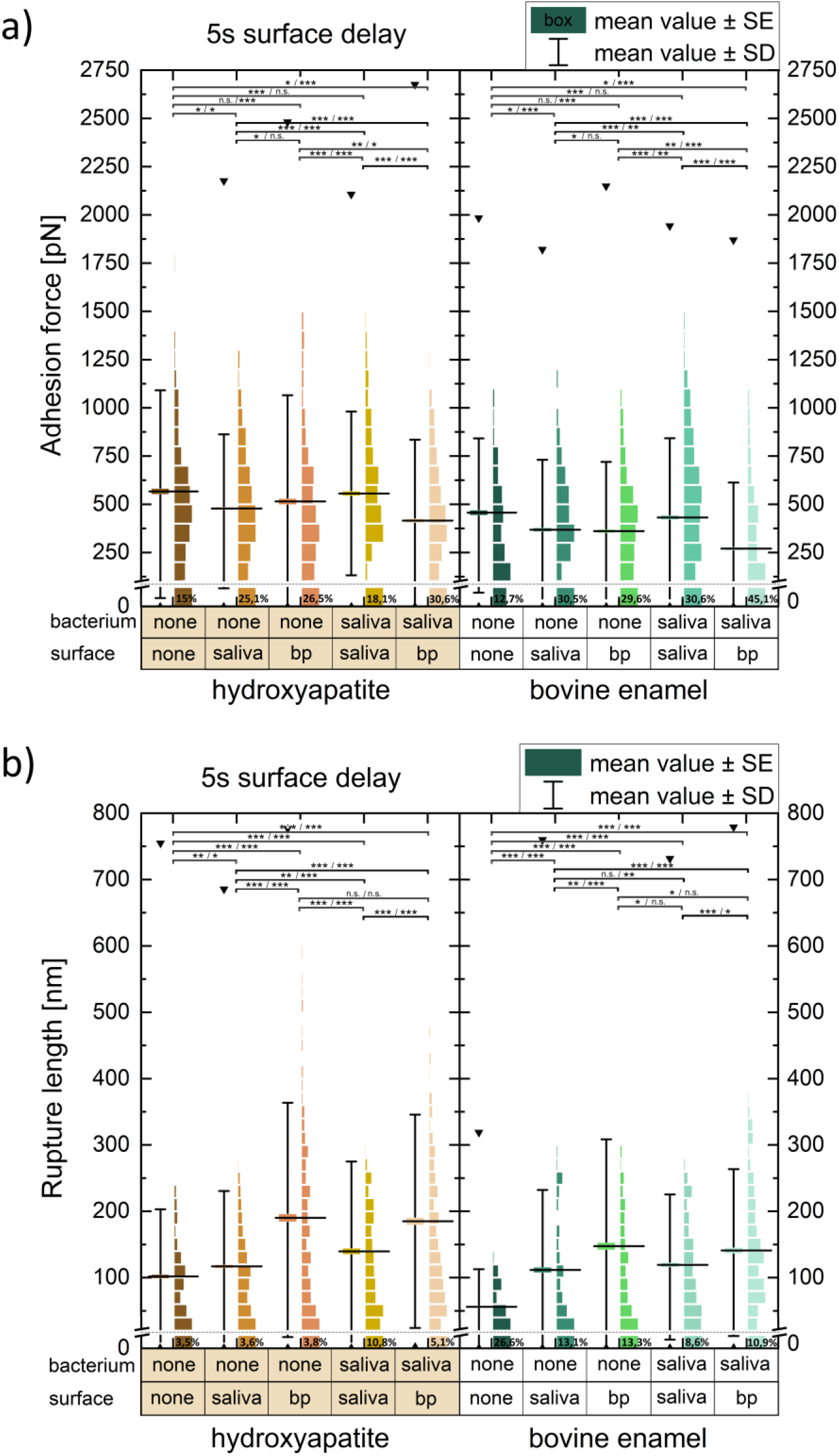
Single-cell force spectroscopy results of 146 *S. aureus* cells on HAp and enamel under all tested conditions. a) Adhesion forces and b) rupture lengths collected from 16-20 bacteria per column with 50 force-distance curves each. The mean value (including all values) is given by a black horizontal line. The boxplot’s box provides the standard error around the mean value and the whiskers the standard deviation. The black triangles represent the maximum values measured.

Throughout all measurements on both surfaces the highest mean adhesion force is generally found on uncoated surfaces with untreated bacteria (see fig. 3). Thus, any of the bodily fluid combinations tested in this paper renders the adhesion of a bacterium less likely than the uncoated state. The rupture length is generally lowest for the combination of uncoated bacteria and surfaces, because only the adhesive macromolecules of the bacterium itself are involved (see fig. 3b). The rupture length of combinations where BP macromolecules were involved are highest, and the saliva-coated surfaces (with and without saliva-treated bacteria) place in between. It is possible that additional hydrogen and ionic bonds can be established on a conditioning film formed on the substratum surface [59]. Whether these are formed or are stronger than the binding of the conditioning film to the surface, depends on the bacterial species, the binding partners involved and the subsurface [4, 29, 38, 60]. For instance, we observe higher rupture length for BP coatings on the surface but no increase in adhesion force compared to the uncoated state. We cannot be certain whether we stretch a weak binding or detach macromolecules from the conditioning films on the surface. If we assume that we detach macromolecules and the binding is so strong that these macromolecules remain bound to the bacterium, we should see a change in adhesion forces between the first force-distance curves and the last ones of each bacterium. We have, however, no indication that the bacteria picked up or detached any macromolecules from the conditioning films which influence the adhesion force, as the first force-distance curve of all bacteria observed can never be considered discordant (see fig. 1b).

Saliva-treated *S. aureus* on BP coated surfaces displayed the lowest adhesion forces in this study. For this condition, the number of observations within the < 100 pN bin was highest among all combinations (see fig. 3a) and the incidence without any adhesion was at least twice as high as those recorded for the uncoated states on each surface, making this the bodily fluid combination with the lowest chance of bacteria adhering at all. Compared to the uncoated states, the mean adhesion force decreases by 27 % (HAp) and 40 % (enamel) for saliva-treated *S. aureus* on BP coated surfaces (see fig. 3a). For practical applications, this finding suggests that HAp-based material covered by BP is likely to reduce the adhesion of planktonic *S. aureus* cells originating from the oral environment, and that a lower force is needed to detach them from the surface. Along the same line is the finding of Gunaratnam et al. who showed that strains of *S. aureus* incubated in BP adhere less to BP covered catheters than bacteria that were not incubated in BP [36].

Notably, the mean adhesion forces of saliva-incubated bacteria on saliva-coated surfaces were just as high as on uncoated surfaces with uncoated bacteria, while only covering the surface with saliva-derived factors decreased the adhesion force. Even though the number of observations within the < 100 pN bin was higher for the double saliva coated (both surface and bacterium) combination, the distribution of adhesion force values above 100 pN for both HAp and enamel was indistinguishable from the ones without any coating (see fig. 3a). The mean, standard error, and deviation in adhesion force of both uncoated and double saliva coated combination were in the same range, while the rupture lengths were higher for double saliva coated combinations, suggesting molecular interactions between salivary factors deposited on the bacterial cell wall and the substratum surface.

When the same bodily liquid (i.e., saliva incubation of bacterium and surface) was used, adhesion forces on both surfaces were in the same range compared to the uncoated state. Different bodily liquids (saliva on bacterium and BP-coating on surface), however, clearly decreased the maximum adhesion forces observed between bacterial cell and substratum. As the blood plasma-coated surfaces were less attractive for saliva-treated bacteria than the saliva treated surfaces, we conclude that a BP conditioning film is more effective in preventing colonization by *S. aureus* than the formation of the physiological salivary pellicle. Similar findings of reduced adhesion of the same and other bacteria, such as oral bacteria, on surfaces such as glass, polystyrene, elastane and polyurethane covered with BP macromolecules further support our proposition [14, 36, 61, 62].

In the case of untreated bacteria, it did not matter for the adhesion forces measured whether enamel and HAp were coated with saliva or BP. Even though the rupture lengths varied, the adhesion force distributions were indistinguishable between the two bodily fluids treated surfaces (see fig. 3a). The percentage of forces measured below 100 pN was also almost identical on each surface and more than 10 % higher than the value for the uncoated surfaces. The resulting overall decreases in mean adhesion force were 12 to 20 %.

Notably, for all combinations of bodily fluids, *S. aureus* cells adhered weaker on enamel than on HAp, although both surfaces exhibited a comparable roughness, similar advancing water contact angles, and consist mainly (97 %) of the same chemical component [43]. Yet, more high-force values were measured, when probing on HAp pretreated with both types of bodily fluids than on enamel surfaces treated in the same way (see fig. 3a). Overall, fewer high-force and high rupture length values were measured on enamel compared to HAp under all tested combinations (see fig. 3a), leading to a 20 to 40 % decrease in both values. These findings demonstrate that adhesion studies conducted with HAp may not necessarily mirror the adhesion forces values that might be seen with the same bacterium on natural enamel. However, the influence of the conditioning films on each surface is remarkably similar. HAp samples therefore still have their value as substitute for natural enamel in dental research, as they provide a reproducible and consistent surface chemistry, which allows controlled alterations in roughness or fluoride content [43, 63, 64]. The development of natural enamel, in contrast, is a highly complex process, which is among others influenced by the individual organic content of enamel and external factors [65], which may lead to larger variations. The non-hydroxyapatite parts of enamel, the ionic substitutes in the mineral component, and/or the crystal orientation, however, seem to make the difference to reach the lowest force values measured in this study. The exact causes for this are a subject for further investigations.

## Conclusion

We compared adhesion of *S. aureus* on HAp to natural enamel, which were both morphologically smooth on a nanometer scale. We showed that regardless of which bodily fluid was used to treat the surfaces, the adhesion force of *S. aureus* decreased significantly in its presence. The lowest adhesion strength was always achieved on BP coated surfaces with saliva incubated bacterial cells. Therefore, it can be concluded that a BP coating best prevents the adhesion of planktonic *S. aureus* cells coming from the oral cavity and also facilitates their removal by providing the lowest forces required for detachment. Overall, the differences observed between HAp and bovine enamel seen in our study are comparably small. Thus, HAp confirmed its value as a surrogate for natural enamel in dental research because HAp pellets provide a reproducible and consistent surface chemistry, which allows controlled alterations such as in roughness or fluoride content. Standardized and well-characterized surfaces like HAp are an essential prerequisite for systematic experimental research on factors influencing bacterial adhesion. Results on, for example, antibacterial coatings or reagents are thus easily comparable and a quantitative correlation can be established by a few measurements on single, well-characterized teeth samples. Performing the same experiments on natural material requires a complete and sometimes very complex characterization for each sample with respect to, for example, roughness, chemical composition of the surface and the material below, crystal domain properties and porosity.

## ASSOCIATED CONTENT

### Supporting Information

0 s surface delay AFM data; SDS-PAGE data of saliva and BP over time; XPS data of enamel and HAp.

## AUTHOR INFORMATION

### Author Contributions

The manuscript was written through contributions of all authors. All authors have given approval to the final version of the manuscript.

## ACKNOWLEDGMENT

The authors thank Philipp Jung and Simone Trautmann for their guidance and advice.

## ABBREVIATIONS

PBS: phosphate buffered saline
AFM: Atomic Force Microscope
XPS: X-ray photoelectron spectroscopy
OCA: optical contact angle
SD: surface delay
BP: blood plasma
HAp: Hydroxyapatite

