## Supporting information for "Hydroxyapatite pellets as versatile model surfaces for systematic studies on enamel"

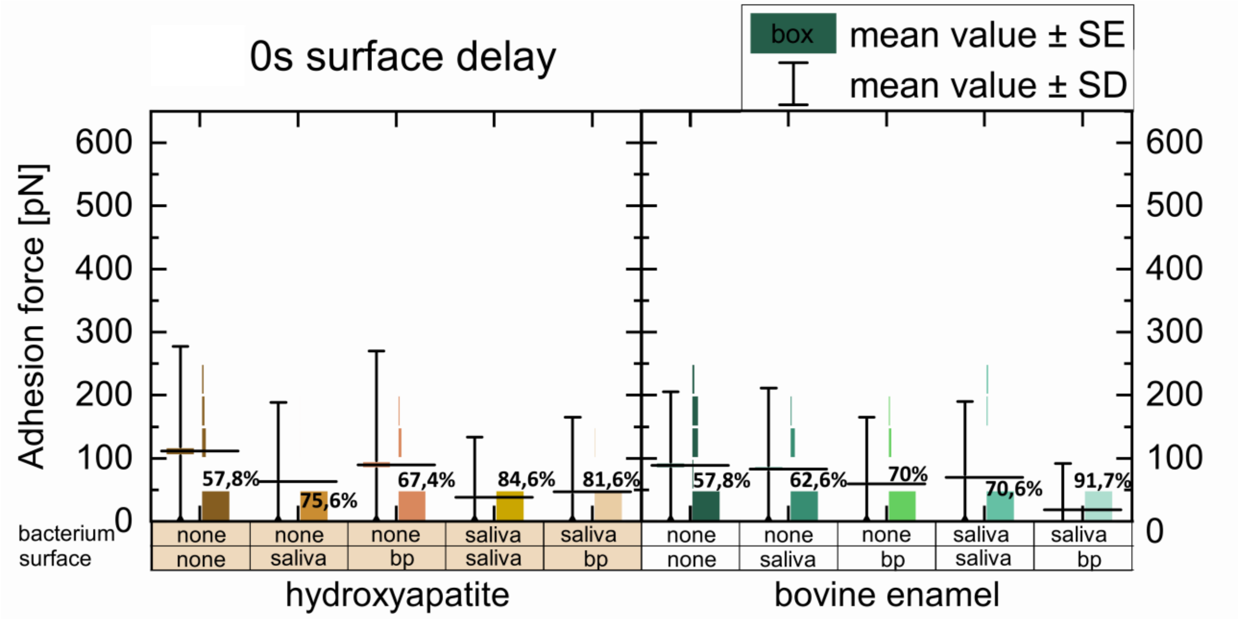


**Figure S1.** Adhesion forces of *S. aureus* cells on titanium implant, HAp and enamel under all tested conditions at 0 s surface delay. The number given in this graph near the 0 - 100 pN area are the percentage of measurement values within the two lowest bins.

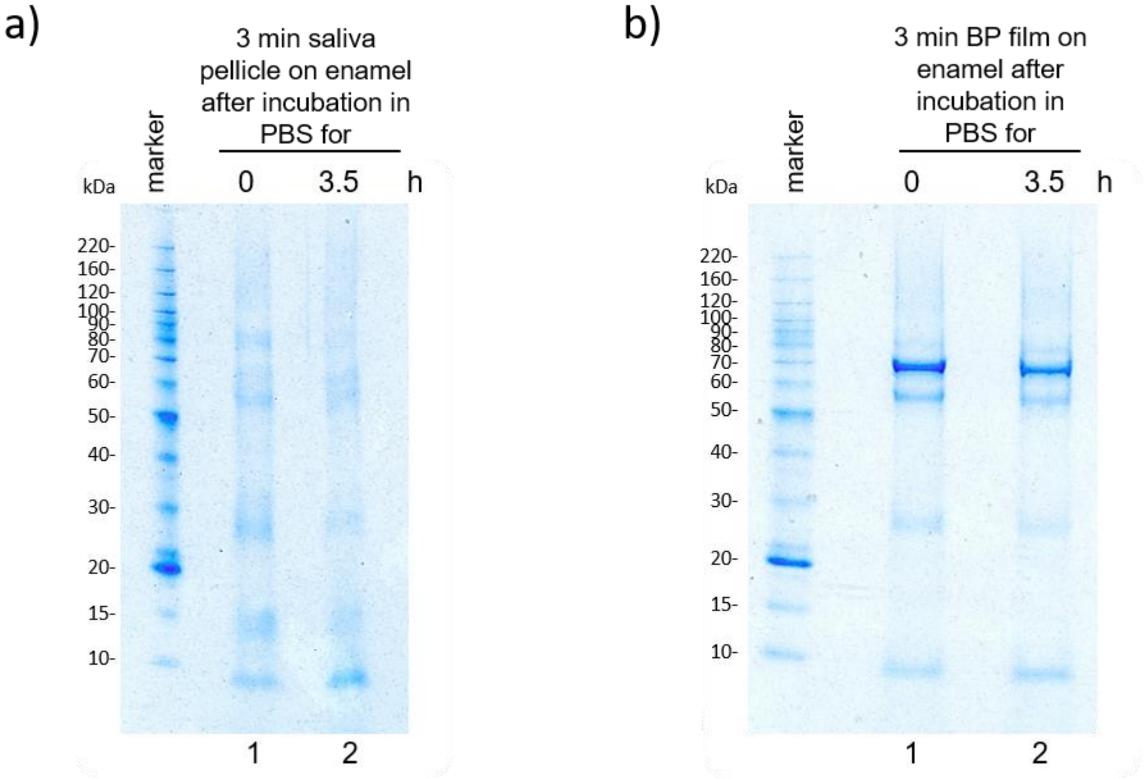


**Figure S2.** Enamel specimens were incubated with saliva a) or blood plasma (BP) b) for 3 min, washed extensively with PBS and either directly eluted (lane 1) or incubated in PBS for 3.5 h (lane 2). After elution of the adsorbed proteins from all samples the eluates were analyzed by LDS-PAGE and Coomassie-staining.

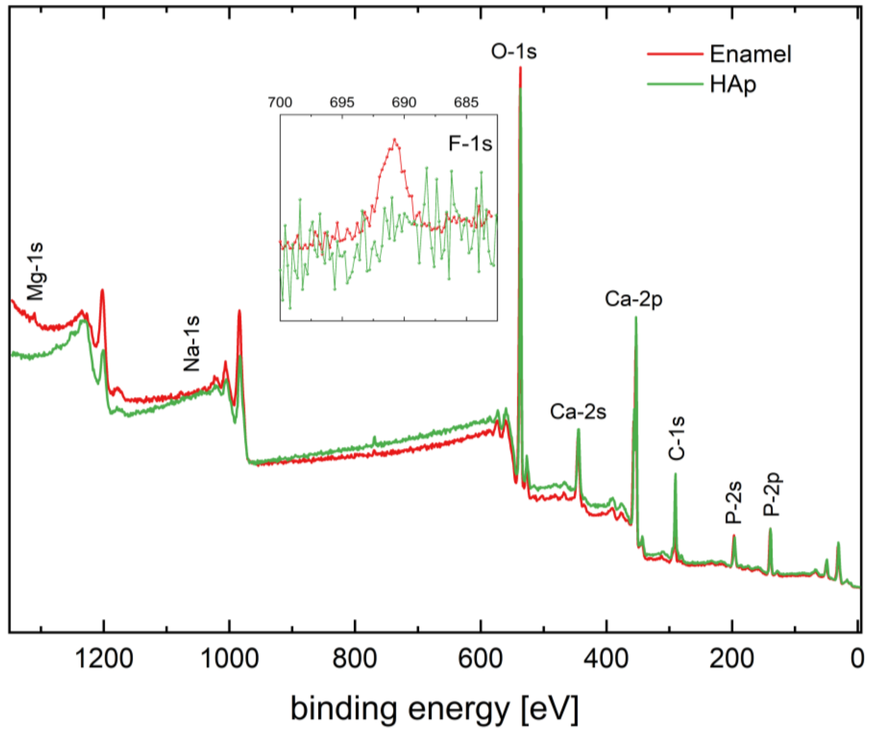


**Figure S3.** XPS survey spectrum of a sintered, non-fluoridated HAp sample in comparison to enamel. HAp contains no Mg, Na and F. The inlay shows the background-corrected signal at the binding energy of fluorine which shows only noise in the HAp signal but a clear peak for enamel.
